# Short-term synaptic depression in multiregional recurrent neural networks accounts for both MMN and P300-like responses and their attentional amplification

**DOI:** 10.1101/2025.11.06.687057

**Authors:** Anthony Strock, Trang-Anh E. Nghiem, Nathan Trouvain, Percy K. Mistry, Xavier Hinaut, Vinod Menon

**Author notes:** Equal Contribution (Anthony Strock, Trang-Anh E. Nghiem). **Address for correspondence:** Anthony Strock, Vinod Menon.

## Abstract

The brain must continuously extract salient cues from an immense sensory stream and represent them across regions. Multiregional responses to salient stimuli, notably mismatch negativity (MMN) in sensory and P300 in frontal areas, are among the best-characterized signals in human electrophysiology, yet the mechanisms relating them are unclear, and whether one mechanism accounts for both is unknown. We develop a hierarchical, multiregional recurrent neural network of sensory and frontal cortex to test whether short-term synaptic depression (STD) can explain both. In sensory regions, STD produces stimulus-specific adaptation, generating MMN-like responses; propagation to frontal regions produces the amplification and delay characteristic of the P300, while enhancing noise robustness, with rare stimuli occupying expanded regions of neural state space. When STD acts on frontal feedback predicting upcoming stimuli, it further amplifies responses to unexpected stimuli, suggesting attention sharpens predictions. Our results bridge synaptic-scale mechanisms and brain-wide representations, with relevance to precision psychiatry.

## Introduction

The human brain is bombarded by an immense stream of sensory data from which it selectively prioritizes stimuli that are rare and behaviorally relevant. The capacity to detect and respond to novel, unexpected, or behaviorally significant stimuli represents one of the most fundamental operations of neural systems, enabling rapid adaptation to changing environments while filtering out irrelevant background information and allocating resources to salient stimuli^1–5^. The failure of this attentional system has profound consequences, contributing to core deficits observed in neuropsychiatric disorders such as schizophrenia, where individuals show impaired ability to distinguish behaviorally-relevant signals from neural and environmental noise^6^. Understanding the neural mechanisms underlying this process is therefore crucial for both basic neuroscience and clinical research.

Event-related potentials (ERPs) have proven invaluable for probing the temporal dynamics of orienting attention in the human brain^7–9^. Among the brain’s electrophysiological signatures, few phenomena are as robust and well-characterized as the neural responses to unexpected stimuli. The discovery of ERPs in response to salient cues represents one of the oldest and most consistently replicated findings in the history of human neuroscience^10–13^, with these responses observed across sensory modalities and experimental paradigms.

The oddball paradigm, where sequences of frequent standard stimuli are occasionally interrupted by rare deviant stimuli, has proven particularly powerful for isolating neural mechanisms of salience detection^14,15^. This simple but powerful experimental design effectively isolates brain responses to unexpected events and has been instrumental in characterizing two key ERP components that represent distinct stages of uncertainty and attention processing^16^.

The mismatch negativity (MMN), typically occurring 150-250 milliseconds post-stimulus, is thought to reflect the brain’s automatic, pre-attentive detection of deviations from established patterns^14^. Originating primarily in sensory cortical areas, the MMN represents a fundamental change-detection mechanism that operates below the threshold of conscious awareness ^17^. Critically, MMN amplitude scales inversely with stimulus probability, providing a direct neural readout of the brain’s internal statistical model of the environment^18^.

The P300 response, peaking approximately 300 milliseconds post-stimulus onset, represents a later, more complex stage of attention processing^19^. Unlike the MMN, the P300 is maximal in frontal and parietal association cortex and is strongly modulated by attention and behavioral relevance^20^. The P300 comprises two functionally distinct subcomponents: an early component elicited by novel unexpected stimuli regardless of their behavioral significance, and a later component, which emerges specifically when stimuli require overt attention and behavioral responses^19^. This temporal and anatomical progression from MMN to P300 is thought to index a hierarchical processing cascade from automatic detection to conscious evaluation^11^.

While MMN and P300 have been extensively studied as biomarkers of cognitive function and are consistently altered in neuropsychiatric conditions^21,22^, their joint underlying neural mechanisms remain poorly understood. The MMN has been the focus of considerable mechanistic and computational investigation, with models attributing it to stimulus-specific adaptation and predictive processing within sensory cortex^23,24^. The P300, by contrast, has received markedly less mechanistic attention, and the processes that transform sensory deviance signals into amplified association cortex responses remain largely unknown. Most fundamentally, unified models explaining how both components arise from shared computational principles have been lacking. This knowledge gap limits our ability to understand how cognitive functions emerge from neural circuits and prevents us from developing principled interventions for disorders characterized by attention processing deficits.

Recent advances in computational neuroscience suggest that short-term synaptic depression (STD), a ubiquitous form of activity-dependent plasticity, may provide a crucial piece of this puzzle^25^. STD causes synapses to weaken dynamically in response to repeated activation, creating a natural mechanism for distinguishing rare from common events. When stimuli are presented frequently, their corresponding synapses become depleted, leading to diminished neural responses. Conversely, synapses transmitting rare stimuli remain active, producing stronger neural responses. This mechanism has been proposed to explain MMN in sensory regions^18^, but its potential role in generating P300 responses and enabling hierarchical attention processing across brain regions remains largely explored.

Several fundamental questions about P300 generation and attention processing remain unresolved. A key unresolved question is whether MMN and P300 can arise from a shared neural mechanism. While STD has been shown to explain MMN in sensory regions^24^, it remains unclear whether the same mechanism can account for P300 in association cortex. The role of feedforward and attentional modulation on P300 dynamics also remains an open question. Another unresolved question is how inter-regional connectivity influences the emergence of P300 signals. While it is well established that P300 responses involve frontal association cortex, the specific role of inter-regional connectivity in shaping these responses is not understood^20^. Although P300 amplitude increases under active conditions that require behavioral response, the underlying neural processes responsible for this enhancement are also not known^19^. Feedback mechanisms from decision-making and attention systems may rely on the same synaptic mechanisms that generate passive responses to oddball stimuli, but this hypothesis has not been computationally tested^26,27^. Evidence accumulation processes may further contribute by sustaining neural representations over time, potentially modulating both the duration and robustness of salient oddball responses^28^.

To test whether MMN and P300 modulation can arise from a single shared neural mechanism, we developed a minimal neural network model of sensory and association cortex. Recurrent neural networks (RNNs) have emerged as powerful tools for bridging explanatory levels in neuroscience, providing a flexible framework for modeling how neural dynamics give rise to cognition and behavior^29–31^. The RNN formalism is particularly powerful because enables systematic exploration of how basic synaptic mechanisms can generate complex neural signatures and explain cognitive phenomena^29–31^. RNN models have been used to demonstrate that STD can reproduce MMN-like responses; however, this model was confined to a single sensory stage and addressed neither the P300 nor behaviorally driven attentional modulation. Recent work has begun extending RNNs beyond individual brain regions^32,33^, and suggested significant potential in RNN development to explore neural mechanisms underlying ERPs and multiregional representations of cognitive processing. Our work advances this frontier by developing the simplest possible hierarchical RNN architectures that model coordinated dynamics across sensory and association cortices, incorporating simple, unified synaptic plasticity mechanisms, and systematically characterizing the representational geometry and robustness that emerges from these dynamics.

We had three fundamental goals which address these longstanding unresolved questions. Our first goal was to validate STD as a sufficient minimal mechanism for sensory-level deviance detection. We systematically tested whether short-term synaptic depression can account for MMN properties observed in sensory regions. While STD has been implicated in sensory adaptation, no previous work has demonstrated its sufficiency for explaining MMN’s probability scaling (inverse relationship with deviant frequency), and temporal sensitivity (dependence on interstimulus intervals). By implementing STD in a minimal sensory network and comparing model predictions to empirical MMN properties, we establish whether this ubiquitous synaptic mechanism provides a complete account of automatic salience detection.

Our second goal was to extend this model and probe mechanisms underlying the P300 and elucidate how multiregional RNNs transform local change detection into enhanced attention signals across sensory and association cortex. We investigated whether the propagation of sensory signals to association regions through feedforward connectivity can explain the transformation from brief, localized MMN responses to the amplified, sustained P300 responses observed in association cortex. We determined the quantitative relationship between inter-regional connectivity and functional amplification, and whether hierarchical processing confers computational advantages such as enhanced noise robustness.

Our third goal was to determine whether passive and active P300 paradigms can be explained by unified mechanisms. We tested the hypothesis that the same STD mechanisms explaining MMN and passive P300 responses can account for attentional enhancement of behaviorally relevant stimuli in active oddball paradigms, which require participants to make an explicit behavioral response^19^. This goal directly addresses the longstanding question of whether passive and active P300 components reflect fundamentally different neural systems or represent different expressions of shared computational principles.

Across each of these goals, we sought to characterize the representational geometry of MMN and P300 processing. The amplitude and latency measures that dominate ERP research summarize the population response as a scalar at each point in time, leaving unaddressed how stimulus information is actually organized across neurons, and therefore how reliably it can be read out by downstream regions^34,35^. To address this, we investigated how stimulus representations organize in high-dimensional neural state space, with particular focus on: (a) how representational manifolds scale with stimulus probability and frequency, (b) how hierarchical processing transforms the geometry of these manifolds, and (c) whether geometric changes enhance the robustness of stimulus discrimination under noise. This analysis addresses the crucial but understudied question of how information is spatially organized in neural networks and how this organization supports cognitive function, a question that traditional amplitude-based ERP analyses cannot address.

Our findings demonstrate that a parsimonious RNN model incorporating STD with feedforward and feedback connectivity can explain the full spectrum of neural responses to behaviorally salient stimuli observed in both passive and active oddball paradigms. Our findings reveal that remarkably simple principles, STD combined with inter-connected multiregional RNNs, can explain the emergence of complex neural signatures underlying the P300 ERP complex. More broadly, this work demonstrates how computational approaches can illuminate the fundamental principles that govern the transformation of neural activity into cognitive function, providing a template for understanding other complex brain functions.

## Results

### A. RNN modelling of deviancy detection

#### Short-term synaptic depression explains enhanced responses to deviant stimuli

To establish a foundational mechanism for deviancy detection, we first investigated whether STD could account for the enhanced neural responses to rare stimuli observed in sensory regions during oddball paradigms. We modeled a sensory cortical network receiving inputs from two stimulus-specific input units (**Figure 1A**). The model incorporated STD at synapses between input units and the RNN, consisting of 80 excitatory and 20 inhibitory interconnected integrate-and-fire neurons. One of two stimuli (*X* and *Y*) was presented at regular 2-second intervals, with a controlled probability of occurrence ranging from 0.1 to 0.9. Standard stimuli occurred with higher probability than deviant ones, typically 0.8 and 0.2 respectively.

**Figure 1.**
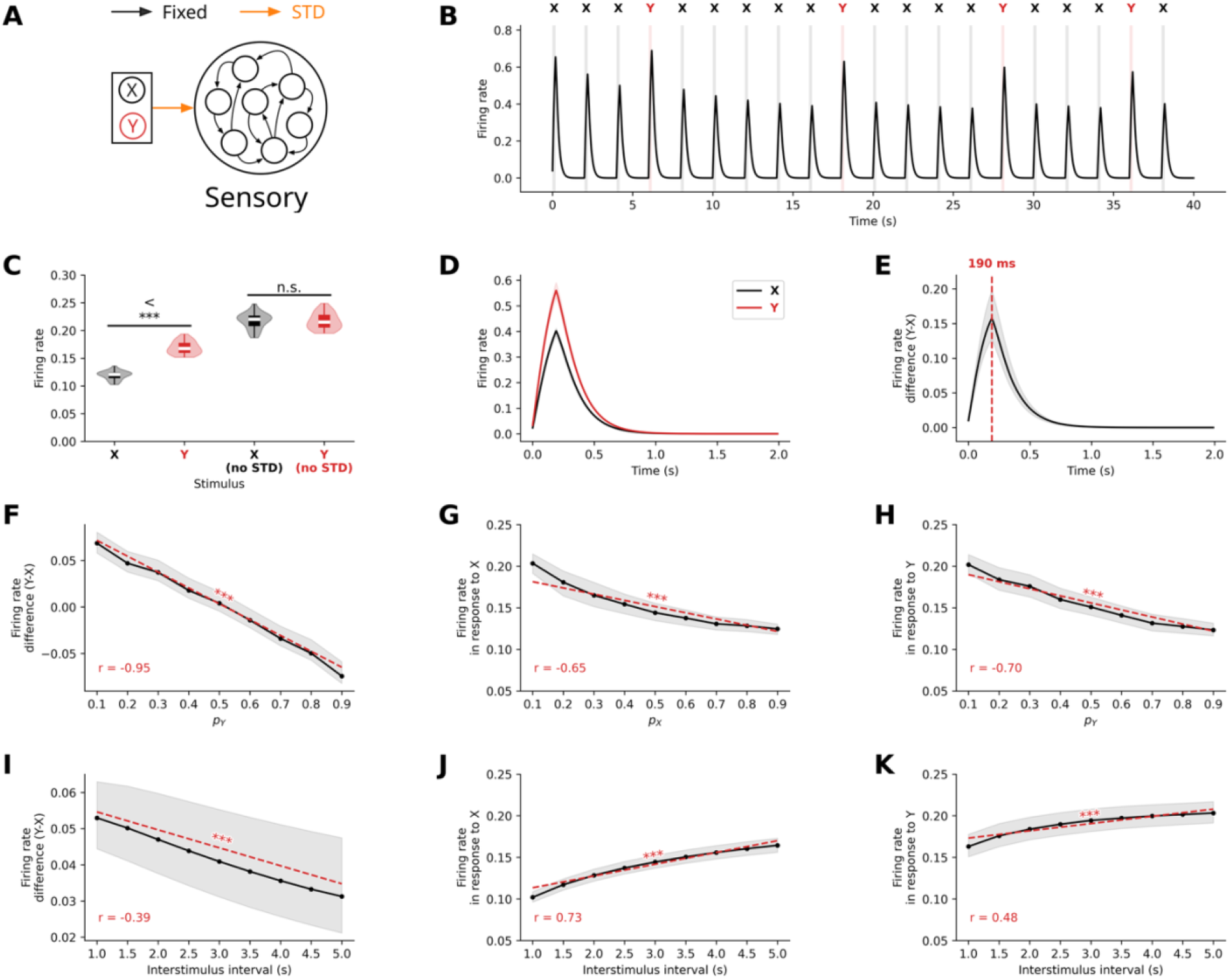
Short-term synaptic depression (STD) accounts for enhanced sensory responses to deviant stimuli in a passive oddball paradigm. **(A)** Schematic of the recurrent neural network model with STD modulating input connection strength. **(B)** Representative time series showing mean firing rate within the sensory network during oddball stimulus presentations. Gray bars indicate stimulus presentations with stimulus identity labeled above (X = standard, Y = deviant). **(C)** Comparison of mean firing rates for standard (X) and deviant (Y) stimuli with and without STD implementation. STD is essential for generating differential responses (*p* < 0.001). **(D)** Population-averaged firing rate responses to standard and deviant stimuli over time, showing enhanced responses to deviant stimuli. **(E)** Time course of the deviant-standard response difference, with peak occurring at 190ms post-stimulus onset (median, red dashed line). **(F)** Mean deviant-standard response difference as a function of deviant stimulus probability, showing inverse relationship. **(G-H)** Individual stimulus responses as functions of their respective probabilities, demonstrating that both stimulus types show probability-dependent response amplitudes. **(I)** Mean deviant-standard response difference as a function of interstimulus interval, showing decay with longer intervals **(J-K)** Individual stimulus responses as functions of interstimulus interval, showing differential increases to standard and deviant stimuli. Data represent median ± interquartile range.

We observed that STD at input synapses systematically modulated neural responses in a probability-dependent manner. Standard stimuli, due to their frequent presentation, induced progressive neurotransmitter depletion at their corresponding synapses, resulting in diminished response amplitudes over time (**Figure 1B**). In contrast, synapses transmitting deviant stimuli had more time to recover between presentations, better preserving their efficacy, and producing stronger neural responses.

The difference in the firing rate of sensory neurons between deviant and standard stimuli was statistically significant (*t* = 18.46, *p* < 10^−1^^6^, **Figure 1C**), but was no longer significant when STD was removed (*t* = −0.50, *p* = 0.96, **Figure 1C**). The peak differentiation consistently occurred at 190 ms post-stimulus onset (**Figure 1D-E**), coinciding roughly with stimulus offset. Moreover, this difference scaled inversely with deviant probability difference (*r* = −0.95, *p* < 10^−1^^35^, **Figure 1F**) and interstimulus interval (*r* = −0.39, *p* < 10^−10^, **Figure 1I**), matching known MMN characteristics.

Further analysis confirmed significant negative correlations between firing rate and stimulus probability for both stimuli (for stimuli *X*, *r* = −0.65, *p* < 10^−3^^23^, **Figure 1G**, for stimuli *Y*, *r* = −0.70, *p* < 10^−3^^23^, **Figures 1H**), demonstrating that single-neuron response inherently encode stimulus probability. This explains why the difference between average responses to stimuli B and A is negatively correlated to *p_Y_* − *p_X_* = 2*p_Y_* − 1, and therefore highly negatively correlated to *p_Y_* (**Figure 1E**).

In contrast, firing rate showed a positive correlation with interstimulus interval for both stimulus types (for stimuli *X*, *r* = 0.73, *p* < 10^−3^^23^, **Figure 1G**, for stimuli *Y*, *r* = 0.48, *p* < 10^−276^, **Figures 1H**), but faster increase for standard stimuli (**Figure 1G**) than for deviant stimuli (**Figures 1H**). Since, the average response for both stimulus types tends toward the overall firing rate when STD is removed, and the response to standard stimuli is lower than that to deviant stimuli at a 2-second inter-stimulus interval, this explains why the difference between the average responses to stimuli *Y* and *X* decreases as the inter-stimulus interval increases.

These results establish that STD at input synapses is sufficient to explain the characteristic properties of MMN responses in sensory cortex, including appropriate latency (150-250ms), inverse scaling with deviant probability, and sensitivity to interstimulus intervals. Moreover, our findings reveal that probability encoding can naturally emerge from this simple synaptic mechanism.

#### Frequency-dependent neural representation geometry emerges from STD

To investigate how STD shapes the distributed representation of stimuli at the population level, we analyzed the high-dimensional patterns of neural activity in the sensory node. Neural activity from all 80 excitatory neurons in the sensory network was recorded during 100 trials of the oddball paradigm, varying either the probability of occurrence of stimuli or the interstimulus interval, both of which affect the frequency of occurrence of stimuli.

We applied two distinct principal component analyses (PCA) to simulated population activity matrices. Activity matrices were of dimension *N* × *T* with *N* = 80 (number of neurons) and *T* = 1800*s* (number of timesteps following first stimulus onset, with 10ms resolution, when varying probability of occurrence) or *T* = 2700*s* (number of timesteps following first stimulus when varying interstimulus interval). PCA of neural population activity revealed distinct representational patterns for stimuli in the oddball paradigm. The temporal trajectories of neural responses formed two non-overlapping semi-filled elliptical patterns in the reduced dimensional space, with one ellipse corresponding to stimulus *X* and the other to stimulus *Y* (**Figure 2A**). This separation indicates that at the population level, the network maintains distinct neural codes for stimuli based on their identity.

**Figure 2.**
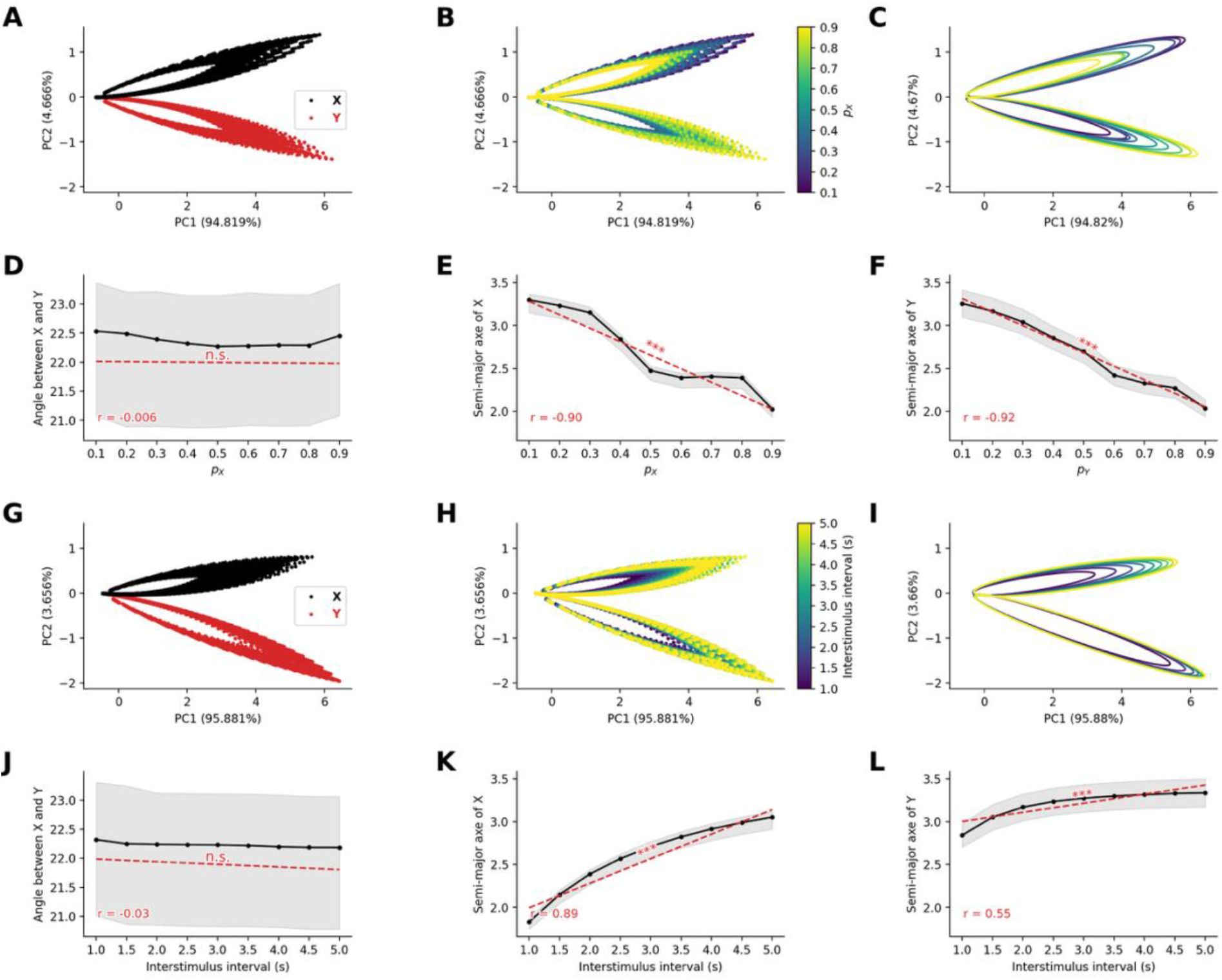
Stable neural representation geometry emerges from short-term synaptic depression in sensory networks. **(A)** Principal component analysis (PCA) of sensory network population responses to standard and deviant stimuli. Neural trajectories form distinct elliptical patterns in PC space, colored by stimulus identity (red = standards A, black = deviants B). **(B)** Principal component analysis of sensory network population responses to standard stimuli, colored by stimulus probability (color scale). **(C)** Fitted ellipses to stimulus representations show probability-dependent geometric properties. **(D)** Angular separation between stimulus ellipses remains constant across probabilities. **(E-F)** Semi-major axes of fitted ellipses decrease with increasing stimulus probability for standard and deviant stimuli, encoding frequency information in representational geometry. **(G-H)** PCA with varying interstimulus intervals at stimulus probabilities of *p_X_* = 0.8 and *p_Y_* = 0.2, showing similar elliptical trajectories colored by (G) stimulus identity and (H) interstimulus interval duration. **(I)** Fitted ellipses for different interstimulus intervals. **(J)** Angular relationships between standard and deviant remain stable across interstimulus intervals. **(K-L)** Semi-major axes increase with longer interstimulus intervals, reflecting enhanced neural differentiation with more recovery time between stimuli. Results demonstrate emergent encoding of stimulus frequency in population-level representational geometry.

When we systematically varied the probability of the stimuli, we observed that the size of the elliptical trajectories was inversely related to stimulus occurrence probability (**Figure 2B**). Specifically, by fitting ellipses to the first second of the last 50 trials (**Figure 2C**), we observe that while the orientation of the ellipses does not vary significantly with probability of stimuli (*r* = −0.006, *p* = 0.92, **Figure 2D**), the semi-major axes of the ellipse decrease with the probability of occurrence for both stimuli (for stimulus *X*, *r* = −0.90, *p* < 10^−9^^6^, **Figure 2E**, for stimulus *Y*, *r* = −0.92, *p* < 10^−1^^10^, **Figures 2F**).

Similarly, when we systematically varied the interstimulus interval, we observed that the size of the elliptical trajectories was related to the interstimulus interval (**Figure 2H**). Specifically, we observed that while the orientation of the ellipses does not seem to vary with interstimulus interval (*r* = −0.03, *p* = 0.58, **Figure 2J**), the semi-major axes of the ellipse increased with the interstimulus interval for both stimulus type (for standard stimuli *X*, *r* = 0.89, *p* < 10^−9^^4^, **Figure 2K**, for stimuli *Y*, *r* = 0.55, *p* < 10^−2^^1^, **Figures 2L**).

These results demonstrate that STD can naturally give rise to emergent neural representations of stimulus frequency, where rarer events occupy larger representational areas in neural state space. This geometric property of neural representations provides a potential mechanism for the amplified processing of surprising or unexpected events at the neural population level.

### B. RNN modeling of passive oddball task

#### Inter-regional connectivity produces amplified P300-like responses in frontal regions

Having established the mechanism for sensory-level deviance detection, we next investigated how these signals propagate and are transformed to generate the P300 component, which originates primarily from (non-sensory) association cortex and exhibits distinct temporal characteristics from the MMN. To accomplish this, we extended our model by adding a second RNN representing “frontal” cortical areas (**Figure 3A**). This frontal network received no direct sensory input but was driven by excitatory projections from the sensory network. We systematically varied the strength and density of these inter-regional connections to examine their impact on frontal response dynamics.

**Figure 3.**
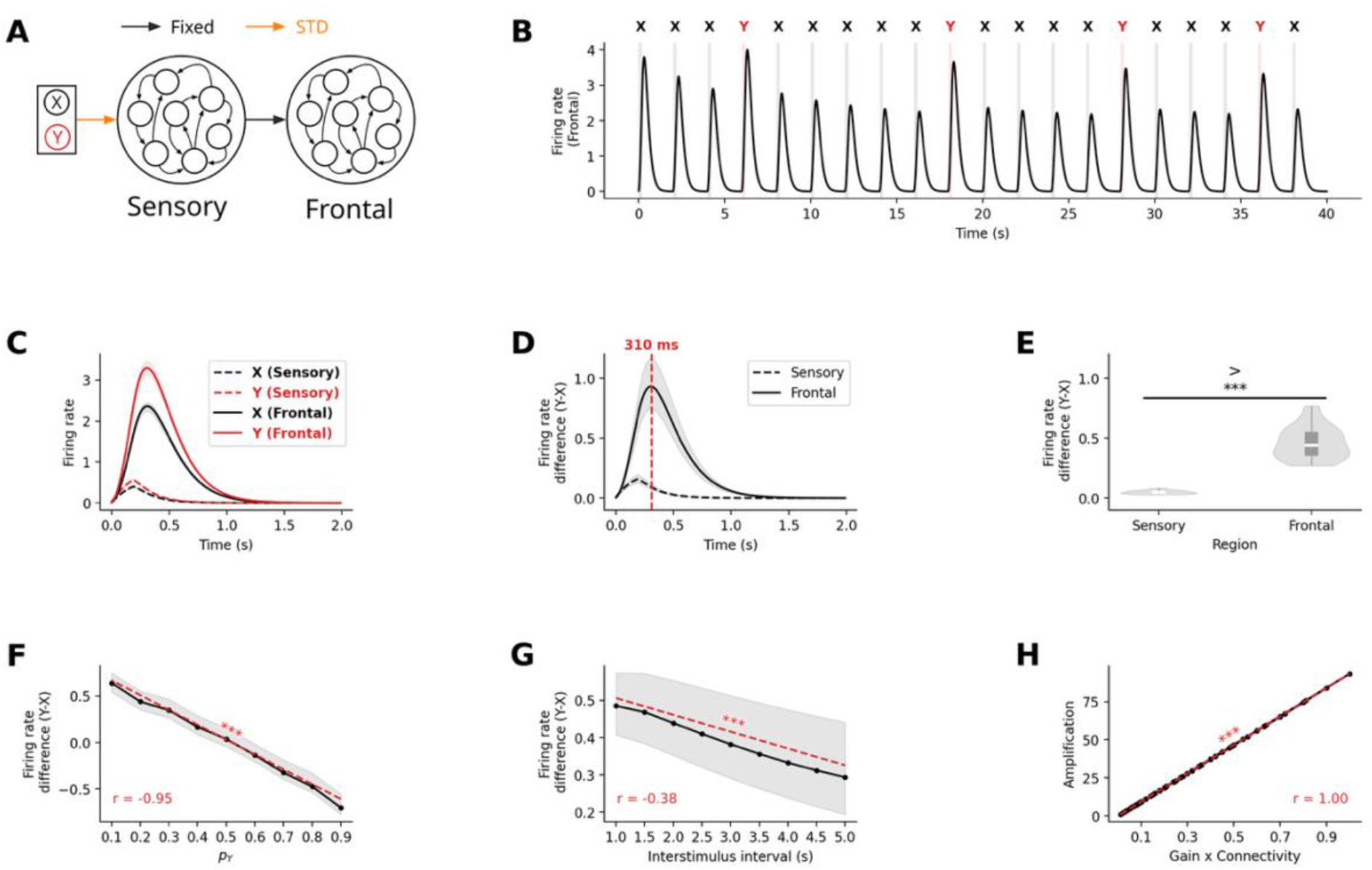
Amplification of deviant-standard response differences from sensory to frontal cortex. **(A)** Extended model architecture with separate sensory and frontal recurrent networks. Frontal network receives excitatory projections from sensory network but no direct stimulus input. **(B)** Mean firing rate within the frontal network during oddball stimulus presentation. **(C)** Direct comparison of firing rates between networks for standard and deviant stimuli, showing delayed, amplified, and prolonged responses compared to sensory responses. **(D)** Temporal evolution of deviant minus standard differences in both networks, with frontal peak delayed by ∼120ms (310ms vs 190ms) and amplified ∼9-fold. **(E)** Quantification of significant response amplification from sensory to frontal regions (*p* < 10^−3^). **(F)** Frontal deviant-standard difference maintains inverse relationship with deviant probability (*p* < 10^−3^). **(G)** Similar inverse relationship with interstimulus interval (*p* < 10^−3^). (**H**) Sensory to frontal amplification ratio scales linearly with the product of inter-regional connection strength and probability (*p* < 10^−3^), revealing the structural basis of functional amplification. This demonstrates that P300-like responses can emerge naturally from hierarchical connectivity without requiring additional mechanisms beyond STD.

With appropriate connectivity parameters (e.g. full density, connection strength *g* = 0.1), neural responses in the frontal network exhibited the hallmark characteristics of P300 signals (**Figure 3B**): they were delayed, amplified, and prolonged compared to their sensory counterparts (**Figure 3C**). Specifically, the peak difference between deviant and standard responses occurred at 310ms in frontal regions compared to 190ms in sensory regions – a 120ms delay (**Figure 3D**). Furthermore, this difference was significantly amplified in frontal areas by a factor of 9.34 ± 0.09 (*t* = 16.59, *p* < 10^−1^^5^, **Figure 3E**).

As with sensory responses, frontal response amplitudes decreased with increasing deviant stimulus probability (*r* = − 0.84, *p* < 10^−40^, **Figure 3F**) and interstimulus interval (*r* = − 0.82, *p* < 10^−6^^6^, **Figure 3G**), consistent with empirical P300 characteristics. Importantly, we found that the amplification ratio between sensory and frontal responses was directly proportional to the product of inter-regional connection strength and probability (*r* = 1.0, *p* < 10^−3^^23^, **Figure 3H**), revealing a simple relationship between structural connectivity and functional amplification.

These results demonstrate that the characteristic properties of P300 response – their delayed timing, enhanced amplitude – can emerge naturally from the propagation of sensory signals through hierarchical brain networks with appropriate connectivity strengths. This mechanism requires no additional computational components beyond STD suggesting a fundamental principle for distributed attentional processing in the brain.

#### P300 responses enhance robustness to noise

Having established that the mechanism underlying P300 component would also cause amplification in frontal regions, we next examined its functional advantage. Specifically, we investigated the downstream processing of salient information by evaluating the conditions of external noise under which the two stimuli could still be discriminated. To this end, we added zero-mean Gaussian noise to the neuronal responses – without altering the underlying neural dynamics – and trained a linear classifier to decode stimulus identity within a 200ms sliding time window.

In the absence of noise, perfect classification accuracy (i.e. 100%) was achieved in both sensory and frontal regions. As the noise standard deviation was systematically increased, classification accuracy degraded rapidly in both the sensory (**Figure 4A&D**) and frontal (**Figure 4B&E**) regions, eventually approaching the performance of a classifier that always predicts the standard stimulus (i.e. 80% given *p_X_* = 0.8). However, we found that as long as the noise standard deviation *σ* remained within reasonable bounds (10^−6^ ≤ *σ* ≤ 1), there existed a time point at which classification accuracy in the frontal network surpassed from 5 to 12% that of the sensory network (*t* ≥ 6.92, *p* < 10^−7^, **Figure 4C**). When considering the entire trial, classification accuracy remained significantly greater in the frontal network than in the sensory network for an even bigger noise standard deviation (10^−6^ ≤ *σ* ≤ 10, *t* ≥ 13.05, *p* < 10^−3^^5^, **Figure 4F**).

**Figure 4.**
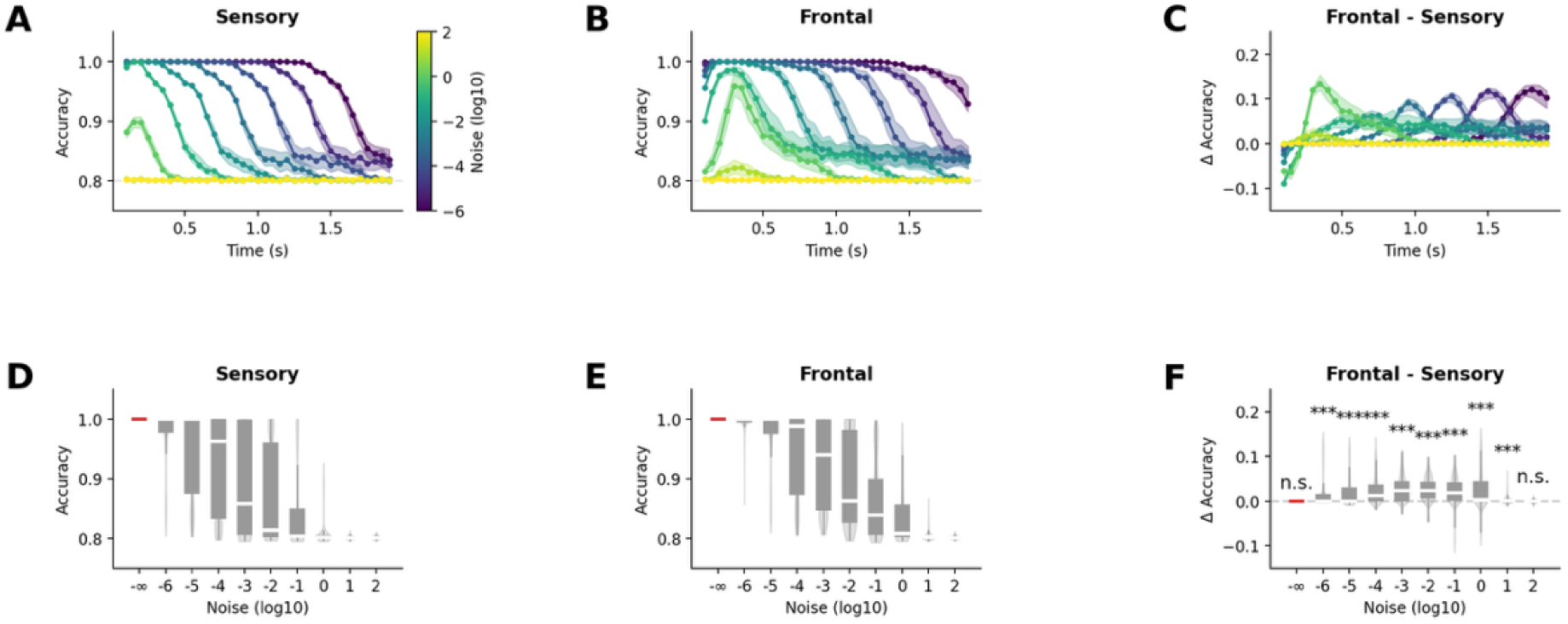
Frontal representations show enhanced robustness to noise compared to sensory regions. **(A-B)** Stimulus decoding accuracy within 200ms sliding windows as a function of post-simulation Gaussian noise amplitude for sensory (A) and frontal (B) networks. Color scale indicates noise standard deviation. **(C)** Difference in decoding accuracy between frontal and sensory networks. **(D-E)** Decoding accuracy across all 200ms windows in 2s trials for varying noise levels in sensory (D) and frontal (E) networks. **(F)** Sustained advantage of frontal over sensory decoding across trial duration, demonstrating functional benefits of response amplification. Red dashed lines indicate constant accuracy levels. Frontal amplification provides significant decoding advantages, supporting the functional role of P300 responses in robust information processing under noisy conditions.

These results demonstrate that the amplification of neural representations in frontal areas can provide a functional advantage by increasing robustness to noise, facilitating more reliable downstream processing of salient information.

#### Frequency-dependent neural representation geometry propagates to frontal regions

To investigate how the frequency-dependent neural representation from the sensory network propagates in the frontal network, we analyzed the high-dimensional patterns of neural activity in the frontal node. Neural activity from all 80 excitatory neurons in the “frontal” node was recorded during 100 trials of the oddball paradigm, while varying either the probability of occurrence of stimuli or the interstimulus interval, both of which affect the frequency of occurrence of stimuli.

We applied two distinct principal component analyses (PCA) to the population activity matrices of dimension *N* × *T*. PCA of neural population activity again revealed distinct representational patterns for stimuli in the oddball paradigm. We observed similar temporal trajectories of neural responses forming two non-overlapping semi-filled elliptical patterns in the reduced dimensional space, with one ellipse corresponding to stimulus *X* and the other to stimulus *Y* (**Figure 5A**). However, in frontal areas, the first component explained even more variance than in sensory areas (∼95% in sensory vs. 99.9% in frontal), and only the third component allowed for stimulus distinction, accounting for just 0.01% of the variance (compared to ∼4% explained by the second component in sensory areas). These results suggest a progressive loss of stimulus-specific information along the processing hierarchy at the population level.

**Figure 5.**
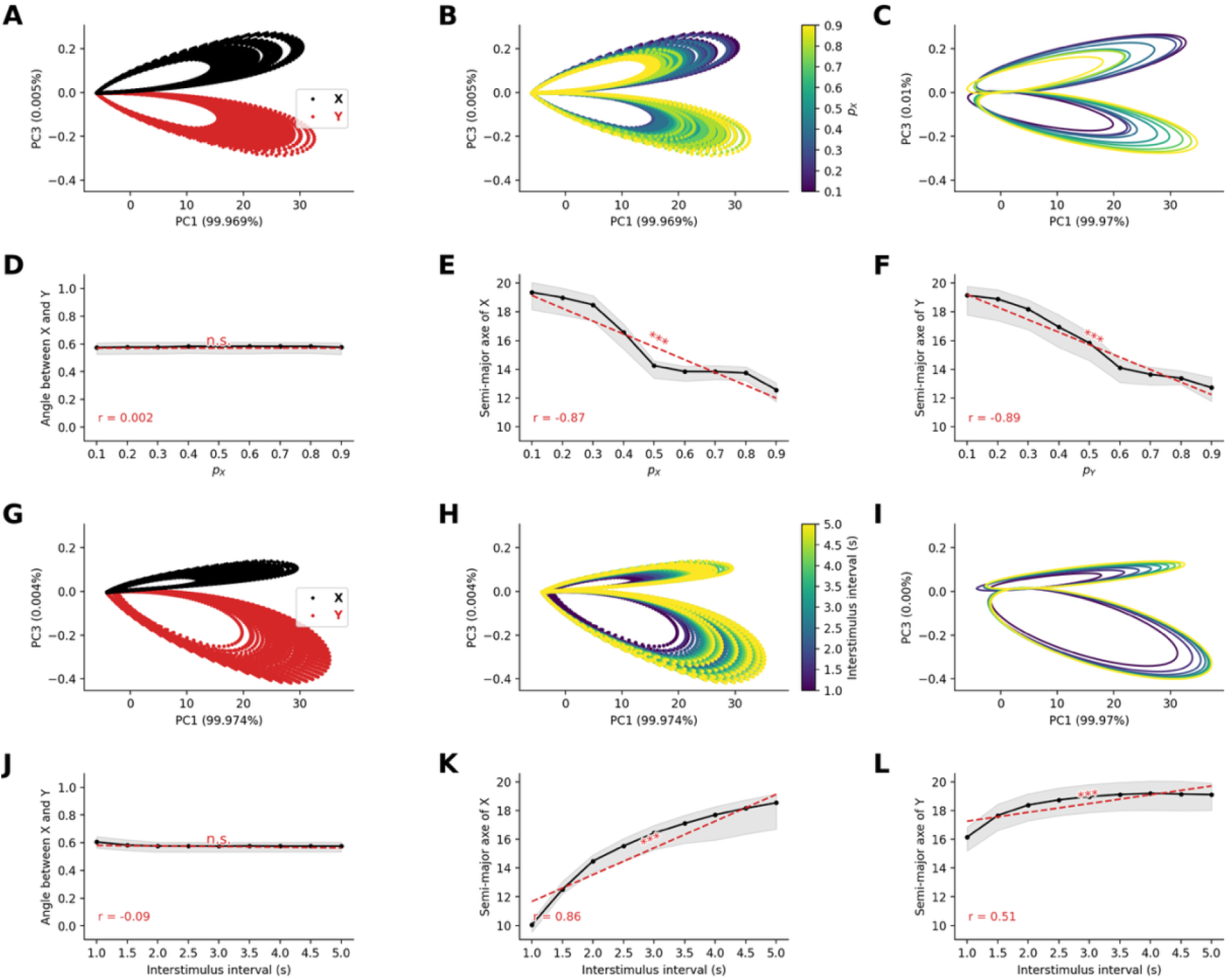
Stable neural representation geometry in frontal networks. **(A-B)** Principal component analysis of frontal network responses with varying stimulus probabilities, showing stability of elliptical representational structure from sensory regions across stimulus intensities (A) and stimulus probabilities (B). **(C)** Ellipse fits to frontal representations preserve probability-dependent geometry. **(D)** Angular relationships remain stable across varying stimulus probabilities. **(E-F)** Semi-major axes of standard (E) and deviant (F) stimuli maintain inverse relationship with probability, demonstrating preserved stimulus frequency encoding. **(G-H)** Analysis with varying interstimulus intervals shows similar preservation of representational structure. **(I)** Ellipse fits across interstimulus intervals. **(J)** Stable angular relationships across interstimulus intervals. **(K-L)** Preserved stimulus interval-dependent scaling of representational size. Results demonstrate that frequency-dependent geometric properties established in sensory regions through STD are maintained and amplified in frontal processing, providing the representational basis for probability-sensitive P300 responses.

As in the sensory network, when we systematically varied stimulus probability, we observed that the size of the elliptical trajectories in the frontal network was inversely related to stimulus occurrence probability (**Figure 5B**). Specifically, by fitting ellipses to the first second of the last 50 trials (**Figure 5C**), we observe that while the orientation of the ellipses does not significantly vary with probability of stimuli (*r* = −0.002, *p* = 0.97, **Figure 5D**), the ellipses’ semi-major axis length decrease with the probability of occurrence for both stimuli (for stimulus *X*, *r* = −0.97, *p* < 10^−8^^2^, **Figure 5E**, for stimulus *Y*, *r* = −0.89, *p* < 10^−9^^3^, **Figures 5F**).

When systematically varied the interstimulus interval, we observed that the size of the elliptical trajectories in the frontal network was related to the interstimulus interval (**Figure 5H**). Specifically, by fitting ellipses to the first second of the last 50 trials (**Figure 5I**), we observed that while the orientation of the ellipses does not seem to vary with interstimulus interval (*r* = −0.09, *p* = 0.13, **Figure 5J**), the semi-major axes of the ellipse increased with the interstimulus interval for both stimulus type (for standard stimuli *X*, *r* = 0.86, *p* < 10^−80^, **Figure 5K**, for stimuli *Y*, *r* = 0.51, *p* < 10^−1^^8^, **Figures 5L**).

These results demonstrate that the frequency-dependent neural representation that emerges in sensory network can naturally be transmitted to the frontal network and explains the observed frequency dependency of P300 amplitude.

### C. RNN modelling of active oddball task

#### Attentional modulation drives enhanced neural responses in active oddball tasks

Having established that STD and inter-regional connectivity can account for the passive oddball P300 response, we next investigated whether the same mechanisms could be extended to explain the enhancement of neural responses in the active task paradigm. Empirical studies consistently show that P300 responses are further enhanced when participants actively respond to target stimuli compared to when they passively observe the same stimuli. This phenomenon, reflected in the P3b ERP component – has been linked to attentional modulation and behavioral relevance. We hypothesized that attentional modulation could account for this enhancement while maintaining the same core STD mechanism that explained passive oddball responses.

To model the active oddball paradigm, we extended our network architecture by adding two readout units connected to the frontal network (**Figure 6A**), which were trained to discriminate between standard and deviant stimuli throughout the whole duration of a trial. These units represented the two possible behavioral responses: producing a motor output (in response to target deviant stimuli) or withholding response (for standard stimuli). The feedforward connections from the frontal network to these readout units were trained using FORCE learning^36^ to ensure that the appropriate unit activated following each stimulus type. Critically, we implemented feedback connections from these readout units back to the frontal network, with these connections also subject to STD. This created an attentional modulation where behavior-related signals could modulate frontal activity patterns (**Figure 6B**).

**Figure 6.**
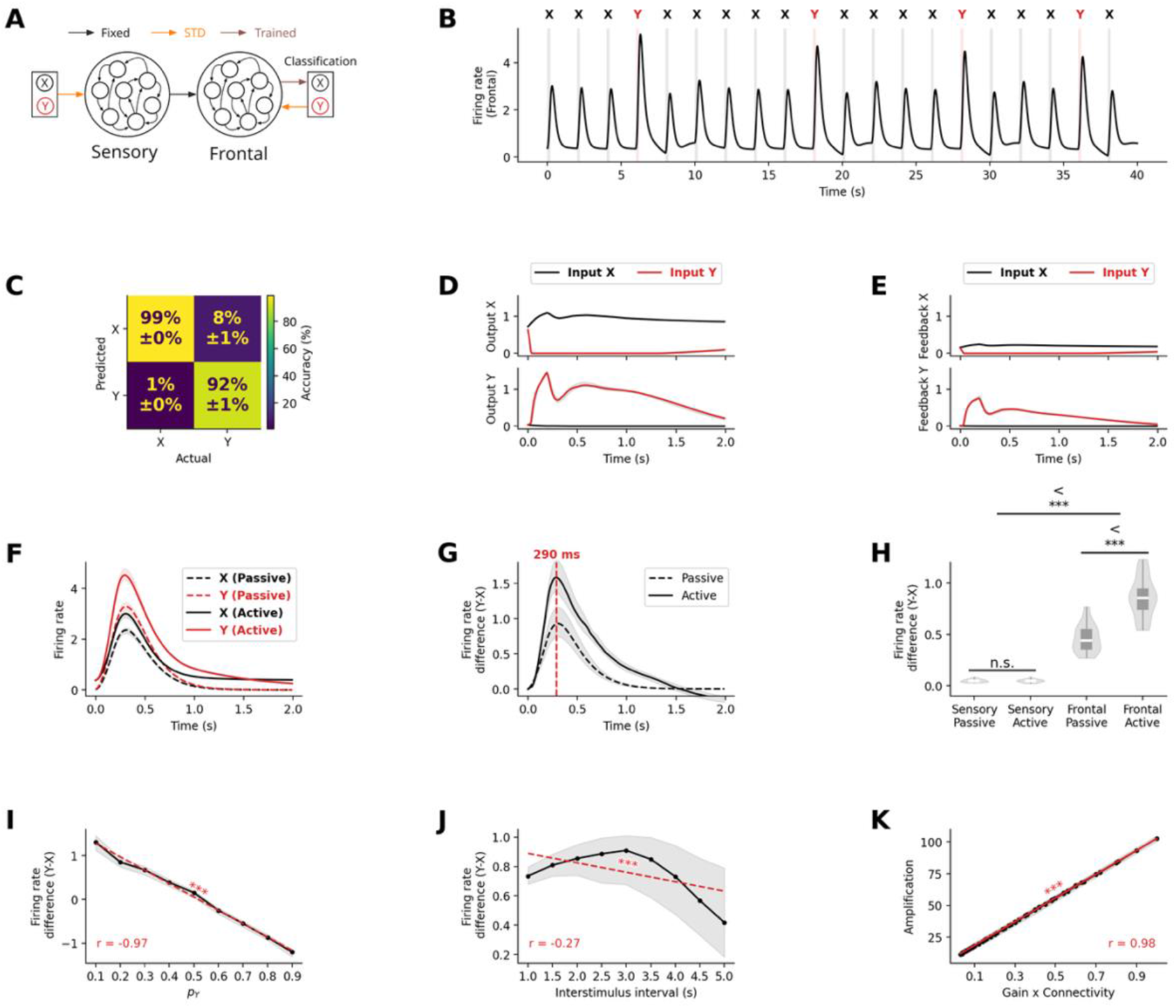
Attentional modulation via feedback STD from decoder enhances frontal network neural response in active oddball paradigms. **(A)** Extended model architecture for active oddball task, incorporating trained readout units with feedback connections subject to STD. **(B)** Frontal network activity during active paradigm shows similar temporal dynamics with additional feedback modulation. **(C)** Confusion matrix demonstrating high classification accuracy (99% for standard stimuli X, 92% for deviant stimuli Y). **(D)** Frontal population responses show differential activations to deviant and standard stimuli. **(E)** Effective feedback strength after STD, showing differential modulation of deviant and standard stimuli. **(F)** Comparison of neural responses in frontal network in active vs passive oddball conditions, demonstrating hierarchical amplification with additional feedback enhancement. **(G)** Deviant minus standard neural responses peak at ∼300ms in frontal network, consistent with P3b timing. **(H)** Significant enhancement in frontal (*p* < 10^−3^) but not sensory (*p* = 0.64) network during active compared to passive oddball conditions. **(I)** Deviant minus standard stimulus neural response with varying probability of deviant stimulus presentation. **(J)** Deviant minus standard stimulus neural response across different inter-stimulus intensities. **(K)** Firing rate response amplification ratio between the frontal and sensory network as a function of the product between synaptic gain and connection probability between networks, showing that linear relationship between connectivity parameters and amplification is maintained during the active oddball task. Results demonstrate that the same STD mechanism explains both passive MMN/P300 responses and active P3b enhancement through differential feedback modulation

After the training phase, the RNN model successfully reproduced the key characteristics of neural responses in active oddball paradigms. As shown in the confusion matrix, classification accuracy was high (**Figure 6C**), demonstrating that the model reliably discriminated between stimulus categories despite the additional processing stage. The readout units successfully discriminated between stimulus types. The non-target unit consistently activated in response to standard stimuli, while the target unit preferentially responded to deviant stimuli, as the units were trained to. Specifically, stimulus *X* (standard) was correctly classified in 99% of time steps, while stimulus *Y* (deviant) was correctly identified 92% of the time, with 8% of instances misclassified as stimulus A (**Figure 6C**).

We found that the temporal activation profiles of the readout units (**Figure 6D**) showed clear differentiation between stimulus types. The non-target unit responded strongly to both standard and deviant stimuli at stimulus onset but maintained its response only when standard stimulus was presented and decayed rapidly when deviant stimulus was presented. The target unit responded strongly only to deviant stimuli and decayed slowly throughout the trial. This differential activation pattern led to distinct feedback effects on the frontal network. When deviant target stimuli activated the target readout unit, the feedback connections enhanced firing rates within the frontal network, producing an amplified response profile (**Figure 6E**). Conversely, when standard stimuli activated the non-target readout unit, the feedback effect was attenuated due to synaptic depression at the more frequently used feedback connections. This resulted in a significantly larger deviant-standard difference in firing rates compared to the passive condition without feedback. In frontal networks, we observe similar profile in the firing rate response to both standard and deviant stimulus, with amplified response in the active condition (**Figure 6F**. The peak difference between responses to deviant and standard stimuli occurred at approximately 300ms (**Figure 6G**), aligning remarkably well with the typical latency of the P3b component observed in human EEG studies. This suggests that the additional processing time required for evidence accumulation naturally accounts for the temporal characteristics of the P3b response.

Importantly, although the difference in firing rate between deviant and standard stimuli did not differ significantly between passive and active condition in the sensory network (*t* = 0.47, *p* = 0.64), in frontal network, the difference was higher in the active condition than in the passive condition (*t* = 8.95, *p* < 10^−11^, **Figure 6H**).

This demonstrates that feedback from attentional mechanisms can enhance amplification of responses to salient stimuli, providing a possible mechanistic explanation for the increased amplitude of P3b responses to behaviorally relevant stimuli.

#### Attentional modulation preserves robustness to noise in frontal region

Next, we investigated how attentional modulation affects the resilience to noise on downstream processing of salient information in the frontal network. As before, we evaluated the conditions of external noise under which the two stimuli could still be discriminated by adding a zero-mean Gaussian noise to the neuronal responses – without altering the underlying neural dynamics – and trained a linear classifier to decode stimulus identity within a 200ms sliding time window.

As the noise standard deviation increased, classification accuracy declined rapidly in both the passive (**Figure 7A&D**) and active (**Figure 7B&E**) conditions, eventually converging toward the performance of a classifier that consistently predicts the standard stimulus (i.e. 80% given *p_A_* = 0.8). We observed that when the noise amplitude remained within reasonable bounds (10^−5^ ≤ *σ* ≤ 1), the classification accuracy remained high throughout the whole trial in the active condition showing a clear advantage of active over passive condition (*t* ≥ 7.88, *p* < 10^−8^). This pattern also held when looking across the entire trial: classification from the frontal network in the active condition continued to outperform the passive condition (10^−6^ ≤ *σ* ≤ 10, *t* ≥ 13.72, *p* < 10^−3^^8^, **Figure 7F**).

**Figure 7.**
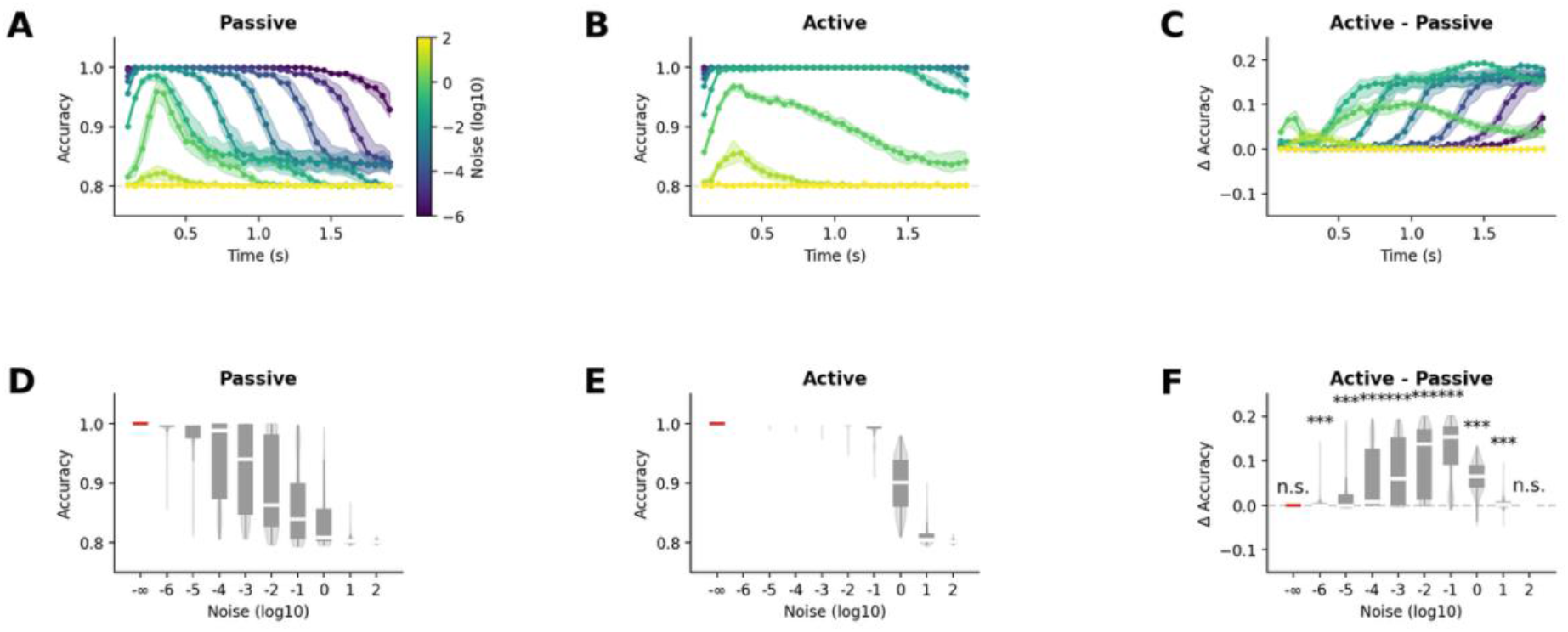
Active oddball paradigm maintains enhanced noise robustness in frontal network. **(A-B)** Decoding accuracy analysis comparing (A) passive and (B) active oddball conditions under varying noise levels. Active condition shows sustained high performance. **(C)** Temporal advantage of active over passive processing, demonstrating sustained functional benefits of amplification with feedback. **(D-E)** Full-trial decoding accuracy for (D) passive and (E) active conditions across noise levels. **(F)** Difference in decoding accuracy between active and passive deviant processing, showing significant improvement across wide range of noise conditions (*p* < 10^−3^). Results confirm that attentional modulation through higher-order feedback connections preserves and enhances the functional advantages of frontal response amplification, supporting robust stimulus discrimination under noisy conditions essential for behavioral responses.

These results demonstrate that attentional modulation can preserve the advantage of the simple amplification observed from sensory to frontal regions in the passive oddball. Consistent with previous results, the amplification of neural representations in frontal areas in active oddball provides a functional benefit by enhancing robustness to noise, thereby supporting more reliable downstream processing of salient information.

#### Frequency-dependent neural representation geometry are preserved by attentional modulation

To investigate how the frequency-dependent neural representation from the sensory network propagates in the frontal network, we analyzed the high-dimensional patterns of neural activity in the frontal node. Neural activity from all 80 excitatory neurons in the frontal network was recorded during 100 trials of the oddball paradigm, varying either the probability of occurrence of stimuli or the interstimulus interval, both of which affect the frequency of occurrence of stimuli.

As before, we applied PCA to the population activity, which again revealed distinct representational patterns for stimuli in the oddball paradigm. We found that the feedback distorted the geometrical elliptical shape (**Figure 8A**). The first component explained less variance in the active than in the passive condition (∼99.9% in passive vs ∼99% in active), and the second component, accounting for ∼1% of the variance, allowed again to discriminate the stimulus identity in the active condition. These results suggest that the feedback from the active condition prevents the loss of stimulus-specific information along the processing hierarchy at the population level.

**Figure 8.**
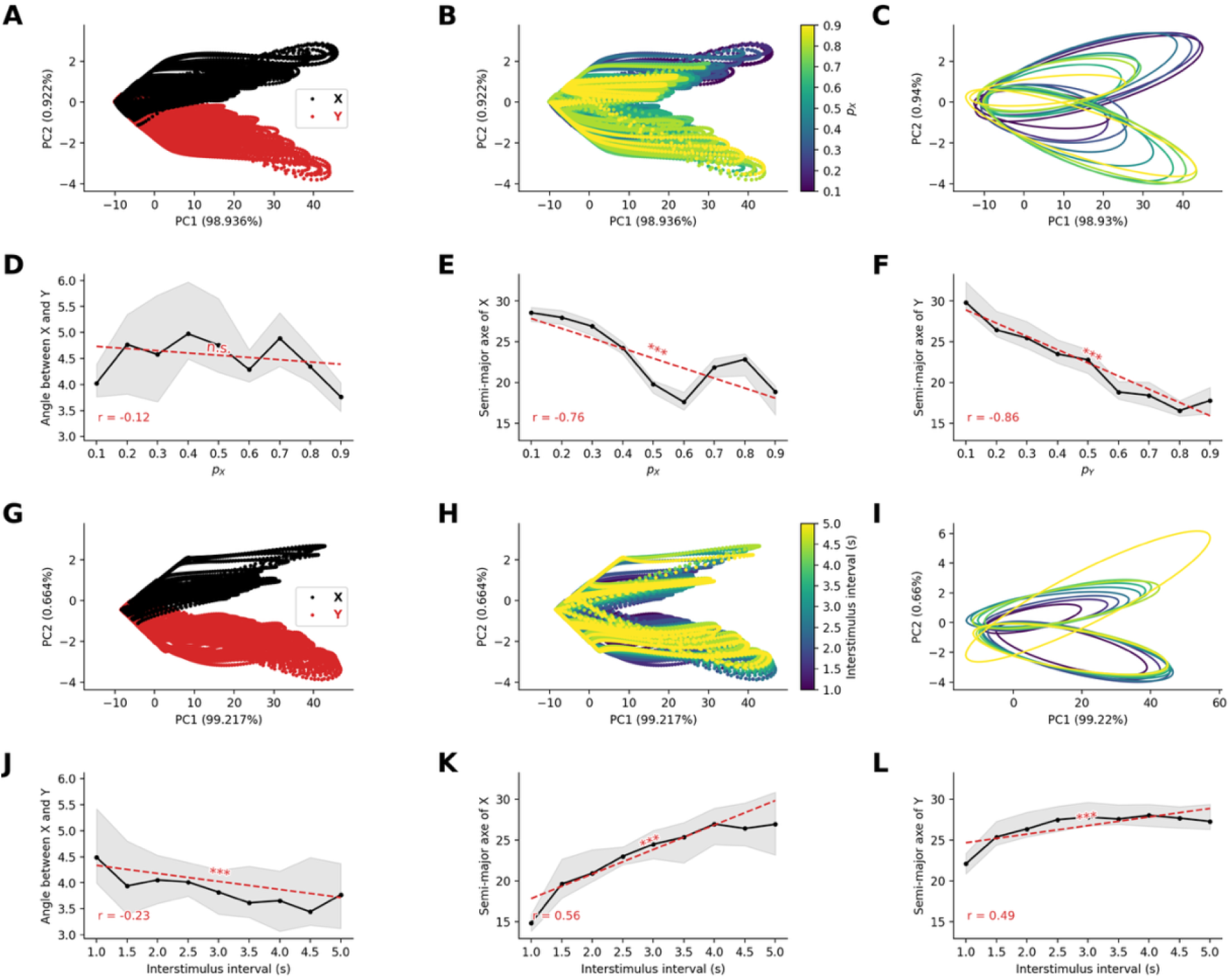
Neural representation geometry is preserved under attentional modulation in active oddball paradigms. **(A-B)** Principal component analysis of frontal responses in active oddball condition with varying stimulus probabilities. Feedback modulation distorts but preserves essential geometric structure, with (A) stimulus identity and (B) probability information maintained. **(C)** Ellipse fits to active condition representations show maintained probability dependencies. **(D)** Angular relationships show slight variation but remain relatively stable. **(E-F)** Semi-major axes preserve inverse relationship with probability. **(G-H)** Analysis with varying interstimulus intervals demonstrates maintained representational structure under attentional modulation. **(I)** Ellipse fits across intervals in active condition. **(J)** Angular relationships show increased sensitivity to interval duration (*p* < 10^−3^). **(K-L)** Preserved scaling relationships between interstimulus intervals and representational geometry. Results demonstrate that attentional feedback preserves the fundamental frequency-encoding properties of neural representations and enhance stimulus-specific information processing, providing a mechanistic basis for the enhanced responses to behaviorally relevant stimuli.

Similar to the active condition, when we systematically varied stimulus probability, we observed that the size of the elliptical contour of the representation in the frontal network was inversely related to stimulus occurrence probability (**Figure 8B**). Specifically, by fitting ellipses to the convex hull of the last 50 trials (**Figure 8C**), we observe that while the orientation of the ellipses does not seem to vary with probability of stimuli (*r* = −0.12, *p* = 0.058, **Figure 8D**), the semi-major axis length decrease with the probability of occurrence for both stimulus type (for stimuli *X*, *r* = −0.76, *p* < 10^−5^^2^, **Figure 8E**, for stimuli *Y*, *r* = −0.86, *p* < 10^−8^^1^, **Figures 8F**).

When we systematically varied the interstimulus interval, we observed that the size of the elliptical contour of the representation in the frontal network was related to the interstimulus interval (**Figure 8H**). Specifically, by fitting ellipses to the convex hull of the last 50 trials (**Figure 8I**), we observed that the orientation of the ellipses slightly vary with interstimulus interval (*r* = −0.23, *p* < 10^−3^, **Figure 8J**), the semi-major axes of the ellipse increased with the interstimulus interval for both stimulus type (for standard stimuli *X*, *r* = 0.56, *p* < 10^−2^^3^, **Figure 8K**, for stimuli *Y*, *r* = 0.49, *p* < 10^−1^^7^, **Figures 8L**).

These results demonstrate that the frequency-dependent neural representation that emerges in sensory network and is transmitted to the frontal network, can be maintained in the active condition and explain the observed frequency dependency of P300 amplitude.

## Discussion

We developed minimal multiregional recurrent neural network (RNN) models to determine whether a single synaptic mechanism – short-term synaptic depression (STD) – can account for the family of oddball event-related potentials (ERPs) that has, to date, been explained by separate and largely descriptive theories. Prior computational work has shown that STD produces stimulus-specific adaptation consistent with the MMN in sensory cortex^24,37–40^. What has been missing in prior computational work is a mechanistic account that links these phenomena: an explicit circuit in which the MMN, the passive P300, and the attention-dependent enhancement of the active-task P300 all arise from the same underlying operation.

Our results demonstrate that short-term synaptic depression (STD) combined with multiregional RNN connectivity and feedback loops can account for a wide spectrum of ERP responses across passive and active oddball paradigms, including their modulation by stimulus probability and behavioral relevance on neural response amplitude and duration. Representational and manifold analysis revealed that RNN populations spontaneously organize into probability-dependent geometric manifolds that enhance noise robustness through hierarchical amplification. These mechanisms generate the rich repertoire of neural responses that emerge during deviant “oddball” processing, from the automatic change detection reflected in mismatch negativity to the complex attentional modulations observed in P300 responses. This represents a fundamental shift from models that invoke separate mechanisms for different cognitive functions toward a unified framework based on shared computational principles operating across multiple regions and scales of brain organization, linking cellular mechanisms to system-level electrophysiology and cognition.

### Short-term synaptic depression as a mechanism for sensory-level detection of salient stimuli

We first used RNNs to validate STD as a sufficient mechanism for explaining the complete spectrum of MMN properties in sensory regions, which we then extended to model the P300. We found that STD can account for the emergence of enhanced responses to deviant stimuli during passive oddball paradigms with remarkable quantitative precision. The model reproduced MMN’s characteristic timing (190ms peak), probability scaling (r = -0.95), and temporal sensitivity (r = -0.39), establishing STD as a viable mechanism for explaining pre-attentive change detection.

Our findings represent a significant theoretical advance beyond previous modeling work. While earlier studies suggested STD’s involvement in sensory adaptation^24,37–40^, none demonstrated its sufficiency for explaining a complete range of empirically-observed MMN phenomenon. Our minimal network architecture, comprising only two input units, STD exclusively on input synapses, and RNN excitatory-inhibitory dynamics, proves that complex cognitive signatures can emerge from elementary synaptic mechanisms. This parsimony challenges assumptions about the need for complex mechanisms. While alternative mechanisms have been proposed, including intracellular potassium depletion^41–43^ and lateral inhibition in retinal circuits^44^, our results suggest STD as a more parsimonious model^15^. The robustness of our findings across parameter variations and the precise quantitative match to empirical data support STD as a candidate mechanism underlying automatic deviance detection. This has important implications for understanding evolutionary conservation, STD is ubiquitous across neural systems, suggesting that deviance detection may represent a fundamental computational primitive present in all nervous systems.

A note on polarity: It should be noted that the MMN deflections observed in our RNN model are positive, similar to other previous computational modeling studies^24^. In contrast, empirical MMN manifests as negative deflections in electric field recordings^11–13,27,45^, reflecting the complex geometry of cortical dipole sources and volume conduction^46^. While our model focuses on the computational logic rather than the biophysical details of electric field generation, it provides a foundation for future work incorporating cortical laminar organization and realistic head models to explain the negative polarity. Crucially, the functional significance, enhanced processing of unexpected events, is fully captured by our STD mechanism.

### Short-term synaptic depression and multiregional RNN mechanisms of the passive P300

Our next goal was to extend the sensory processing model of the MMN to investigate a single STD-based mechanism spanning the oddball ERP complex. To address this, we constructed a multi-regional RNN model and examined how local sensory change-detection signals transform into the amplified, sustained P300 responses characteristic of attentional processing in the context of the oddball task^19,47^. We systematically varied the strength and density of feedforward connections between sensory and “frontal” association RNNs and quantified the relationship between inter-regional connectivity and functional amplification.

Our results revealed that hierarchical connectivity produces a striking transformation of neural dynamics. When inter-regional connections were sufficiently strong and dense (full connectivity with synaptic strength *g* = 0.1), the frontal RNN exhibited all hallmark characteristics of empirical P300 responses: delayed timing (310ms peak versus 190ms in sensory regions), dramatic amplification (9.34-fold enhancement of the deviant-standard difference), and extended duration compared to brief sensory responses. Critically, we discovered a near-perfect linear relationship between the product of connection strength and connection probability and the magnitude of functional amplification (r ∼ 1.0). This relationship held across the full range of connectivity parameters we tested, from weak sparse connections that produced attenuation to strong dense connections yielding substantial amplification.

We also observed that hierarchical processing creates systematic temporal delays that emerge from network dynamics rather than built-in delays. The 120ms delay between sensory and frontal peak responses arose naturally from the filtering properties of recurrent networks, with each processing stage acting as both an amplifier and low-pass filter. This emergent temporal structure matched empirical observations and created increasingly slower dynamics at higher levels of the cortical hierarchy, consistent with longer intrinsic timescales recorded in frontal regions^48,49^.

These emergent properties of temporal hierarchy provide new insights into how cortical organization supports cognitive function. Rather than requiring specialized cellular mechanisms to create different timescales across brain regions, our results suggest that hierarchical network structure can generate P300 ERPs observed empirically. Each RNN in the processing hierarchy naturally implements low-pass filtering that progressively slows responses while amplifying relevant signals. This principle may generalize beyond salience processing to explain temporal hierarchy throughout cortex, suggesting that the characteristic timescales of different brain regions reflect their position in processing hierarchies rather than solely intrinsic cellular properties^50–52^.

To further assess the functional significance of hierarchical amplification, we systematically tested noise robustness. We found that frontal networks maintained superior classification performance compared to sensory networks across a wide range of noise conditions (noise standard deviations from 10^-2^ to 10), with the advantage being particularly pronounced for sustained processing across entire trials. This enhanced robustness persisted even as overall performance degraded with increasing noise levels. The noise robustness suggests a particular advantage of hierarchical brain architectures. By progressively enhancing signal-to-noise ratios through amplification, hierarchical processing may help create representations that remain reliably decodable despite the ubiquitous neural variability that characterizes biological systems.

### Unified mechanisms for passive and active-task enhancement of the P300

Next, we tested whether the same STD and RNN mechanisms explaining passive responses could account for P300 enhancement in active behavioral paradigms. In the active task, the model was required to discriminate the two stimuli throughout each trial: two readout units trained on the frontal population reported the deviant (target) and standard (non-target), and these units projected back to the frontal network through feedback connections that were themselves subject to STD. The model performed the discrimination accurately, and the deviant–standard difference in the frontal, but not the sensory, network was larger than in the passive condition, reproducing the empirical enhancement of the P300 when stimuli become behaviorally relevant. Notably, the ∼300ms peak latency emerged naturally from feedback dynamics rather than being imposed, suggesting that evidence accumulation through the feedback loop intrinsically generates the delay characteristic of active processing.

The longstanding distinction between the passively and actively elicited P300 has been interpreted as evidence for separate systems underlying automatic versus controlled processing mechanisms^19,53^. Our results demonstrate instead that both reflect the same STD mechanism operating in different contexts. The key insight is that behavioral feedback creates asymmetric STD effects: frequent non-target feedback becomes depressed while rare target feedback remains strong, producing selective enhancement for behaviorally relevant stimuli.

### MMN and P300 as readouts of a structured representational geometry

The amplitude and latency measures that dominate ERP research collapse the population response to a scalar at each timepoint, leaving unaddressed how stimulus information is organized across neurons and, consequently, how reliably downstream regions can read it out. This is the question representational geometry is suited to answer: a growing body of work establishes that the geometry of population activity, rather than mean rate alone, governs what a linear decoder can extract and how well that readout generalizes across noise and task context^34,35^. We therefore characterized the geometry underlying the MMN and P300 elicited in the oddball task, asking not only how stimuli are encoded but how that encoding is transformed across the cortical hierarchy.

We found that neural populations spontaneously organize stimulus representations into probability-dependent geometric manifolds, with rare events occupying expanded regions of state space. The inverse relationship between stimulus probability and manifold size emerged from STD dynamics, without explicit probability computation, and provides a principled substrate for both pre-attentive and attentive processing that scales with statistical surprise. Critically, this scaling was preserved as activity propagated from sensory to frontal regions. This reframes the MMN and P300 as readouts of a common, structured geometry expressed at successive stages of the hierarchy.

These observations situate the MMN and P300 within the broader effort to understand computation through neural population geometry^34,35^. The organization of activity into probability-dependent manifolds may be a general feature of circuits that must represent statistical structure, with implications extending beyond attention to memory and decision-making, where analogous geometric reorganizations accompany task acquisition and evidence accumulation^34^. More broadly, our results suggest that the value of these ERP components as biomarkers may rest not on their scalar amplitude alone but on the integrity of the underlying representational geometry, a structure that scalar measures only indirectly reflect, and that population-level recordings could assess directly.

### Clinical relevance and implications

Reduced MMN and P300 amplitudes are among the most replicated electrophysiological findings in schizophrenia^21,54–57^, and P300 abnormalities are also reported in autism spectrum disorder²². Because our model parameterizes the oddball response in terms of synaptic adaptation, excitatory–inhibitory recurrent dynamics, and inter-regional connectivity, it provides a computational model useful for probing how specific, biologically interpretable perturbations, altered STD time constants, weakened recurrent excitation or E/I imbalance, or reduced sensory-to-association connectivity, propagate into changes in MMN and P300 amplitude and latency.

### Limitations

Several limitations qualify our conclusions. Our RNN models capture the computational logic of the oddball response but not the negative polarity or biophysical origin of the scalp-recorded MMN/P300, which depend on cortical laminar geometry and volume conduction; relating population activity to measurable electric fields will require biophysical forward modeling. The model includes a single feedforward step, whereas the empirical generators of the P300 are distributed across frontal and temporal-parietal networks^11–13,27,45^; extending the architecture to additional regions and to top-down projections onto sensory cortex would allow the model to address reported interactions between attention and early sensory responses³¹. We omitted the neuromodulatory systems implicated in the P300; in our framework these would act as complementary gain mechanisms alongside STD. Finally, while the model reproduced the increase of P300 latency and amplitude with stimulus parameters, capturing the full range of empirically reported latency effects will likely require additional manipulation of model parameters including synaptic gain and excitatory-inhibitory balance.

## Conclusions

We show that multiregional recurrent neural networks with short-term synaptic depression, a ubiquitous synaptic mechanism, provides a unified computational framework for characterizing both the MMN and P300. Three advances follow. First, a single mechanism, propagated through hierarchical connectivity and task-driven feedback, generates the spectrum of MMN and P300 oddball responses spanning automatic sensory change detection and attention-dependent association cortex amplification, phenomena previously explained by separate, largely descriptive theories. Second, we find that neural populations spontaneously organize stimulus representations into probability-dependent geometric manifolds, in which rare events occupy expanded regions of state space; this coding strategy emerges from the network’s dynamics, without explicit probability computation, and reframes the MMN and P300 as readouts of a structured representation rather than as isolated waveforms. Third, by parameterizing these signals in terms of synaptic adaptation, excitatory–inhibitory dynamics, and inter-regional connectivity, the model yields testable predictions and for interpreting altered ERPs in disorders such as schizophrenia and autism, where MMN and P300 abnormalities are well-documented biomarkers^21,54–57^. Together, these results demonstrate that a parsimonious principle operating across multiple scales of brain organization can explain complex electrophysiological signatures of attention and offer a parsimonious approach for their modeling in clinical contexts.

## Methods

### Stimuli and task ― Oddball paradigm

To generate stimuli as in the oddball paradigm, we randomly draw one of two stimuli *X* and *Y* with probabilities *p_X_* and *p_Y_* = 1 − *p_X_*. When using a fixed stimulus probability, we set *p_X_* = 0.8 and *p_Y_* = 0.2. For conditions involving varying stimulus probabilities, we systematically varied *p_X_* from 0.1 to 0.9 in increments of 0.1. We refer to the stimulus with the lower probability as the deviant and the one with the higher probability as the standard.

Each stimulus was presented 100 times, with each presentation lasting 200ms. In the fixed interstimulus interval condition, stimuli were presented every 2s, resulting in a total session duration of 200s. For conditions involving variable interstimulus intervals, the interval was systematically varied from 1.0 to 5.0s in increments of 0.5s.

In the passive version of the task, models received stimuli and no outputs were decoded from simulated dynamics. In the active version of the task, model received stimuli and a decoder was trained to reproduce the input from simulated dynamics for the whole duration of the trial, thus mimicking a task detecting the deviant stimuli.

## Model

### A. Recurrent network model of sensory responses to stimuli in passive oddball paradigm

To model thalamic inputs sending stimulus-related signals to the sensory cortex, we simulate two input units corresponding to *X* and *Y* stimuli, whose activity levels are described by a one-hot binary 2-dimensional vector *u* i.e. *u* = (1,0) when stimulus *X* is presented and *u* = (0,1) when *Y* is presented.

We model the sensory cortex as a recurrent network of 80 excitatory and 20 inhibitory neurons subject to short-term depression of input synaptic weights, the dynamics of which can be described by:

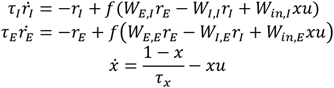

where *r_I_* and *r_E_* represent the firing rate activity of inhibitory and excitatory neurons respectively, *u* represents the 2 excitatory input neurons, one each for stimuli *X* and *Y*, and *x* represents the fraction of neurotransmitters within each input neuron modulating the strength of signal transmission from input units to sensory neurons.

*W_in_*_,*I*_ and *W_in_*_,*E*_ represent the synaptic input weights and are uniformly randomly sampled between 0 and *g_in_* = 1, a positive input gain. *W_E_*_,*I*_, *W_I_*_,*I*_, *W_E_*_,*E*_ and *W_I_*_,*E*_ represent the recurrent synaptic weights and are first uniformly randomly sampled between 0 and 1, then globally multiplied by a scaling factor such that the spectral radius, that is, the eigenvalue product of 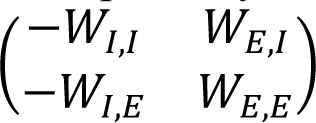 is fixed to *ρ* = 1, and finally scaled individually so that 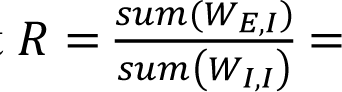 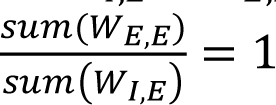.

*τ_I_* = *τ_E_* = 100*ms* represent the time constant of inhibitory and excitatory neurons respectively, *τ_x_* = 20*s* represents the adaptation time constant of the input, and finally *f* represents the non-linear neural response function, and is a rectified linear function (i.e. *f*(*x*) = max(0, *x*)).

To simplify the written representation of the model, we denote the firing rate dynamics as follows:

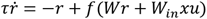

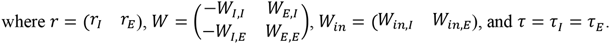

### B. Two-network model of sensory and frontal responses to stimuli in passive oddball paradigm

To model signal transmission from sensory to frontal areas, we simulate the sensory cortex and frontal cortex as two recurrent networks of 80 excitatory and 20 inhibitory neurons each, connected in a feedforward way from the sensory cortex to the frontal cortex, subject to short-term depression only for the input synaptic weights of the sensory cortex. The dynamics can be described by the following:

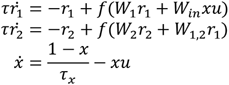

where *r*_1_ and *r*_2_ represent the firing rate of the sensory and frontal cortex respectively, *u* represents input unit activity delivered only to the sensory network, and *x* represents the fraction of neurotransmitters within each input neuron modulating the intensity of the input. *W*_1_ and *W*_2_ represent the recurrent synaptic weights of sensory and frontal cortex respectively, and are sampled in the same was *W* in the previous model. *W*_1,2_ represents the forward synaptic weights from sensory to frontal cortex. Only excitatory to excitatory connections were allowed between cortices. All non-zero weights were randomly drawn from a uniform distribution between 0 and *g* where *g* is a positive gain parameter.

### C. Two-network model of sensory, frontal, and behavioral responses in active oddball paradigm

To investigate the impact of the behavioral relevance of stimuli on frontal responses, we model the behavioral target as an output from the frontal cortex, which is also fed back to the frontal cortex. This feedback is also subject to short-term depression.

To model frontal output detecting the target, we use two output units corresponding to *X* and *Y* stimuli, whose activity levels are described by a one-hot binary 2-dimensional vector *y* i.e. *y* = (1,0) when stimulus *X* is presented and *y* = (0,1) when *Y* is presented.

The overall dynamics of the model can be described by:

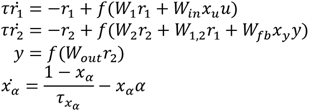

where *r*_1_ and *r*_2_ represent the firing rate of the sensory and frontal cortex respectively, *u* represents the 2 excitatory input neurons, one for each stimuli *X* and *Y*, and y represents 2 excitatory neurons detecting whether the target is present. *x_u_* and *x_y_* represent the fraction of neurotransmitters modulating the intensity of the inputs (within the input neuron) and the target prediction feedback (within the readout neuron) respectively. *W*_1_ and *W*_2_ represent the recurrent synaptic weights of the sensory and frontal cortex respectively, and are sampled similar to *W* in the previous model. *W*_1,2_ represents the forward synaptic weights from sensory to frontal cortex and is defined as in the previous model. *W_out_* represents the synaptic weights allowing the detection of the target from frontal response. *W_fb_* represents the feedback synaptic weights from target detection units to frontal cortex, and are sampled similarly to *W*_1,2_; only target to excitatory neurons connections are allowed and corresponding synaptic weights are randomly uniformly sampled between 0 and *g_fb_* = 2.0, a posititve feedback gain.

*W_out_* is learnt with FORCE learning^36^, a Hebbian learning rule which implements the Recursive Least Squares algorithm^58^. While learning occurs, the feedback that *r*_2_ receives is replaced by the target value for *y* referred as *ỹ*. The overall learning dynamics can be described as:

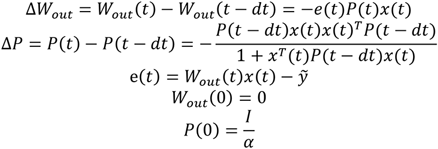

Where *α* = 10^−6^ is a regularization parameter.

### D. Model hyperparameters

We ran 30 instances of every variant of model described above by initializing with different seeds but using the following common hyper-parameters (**Table 1**).

**Table 1.**
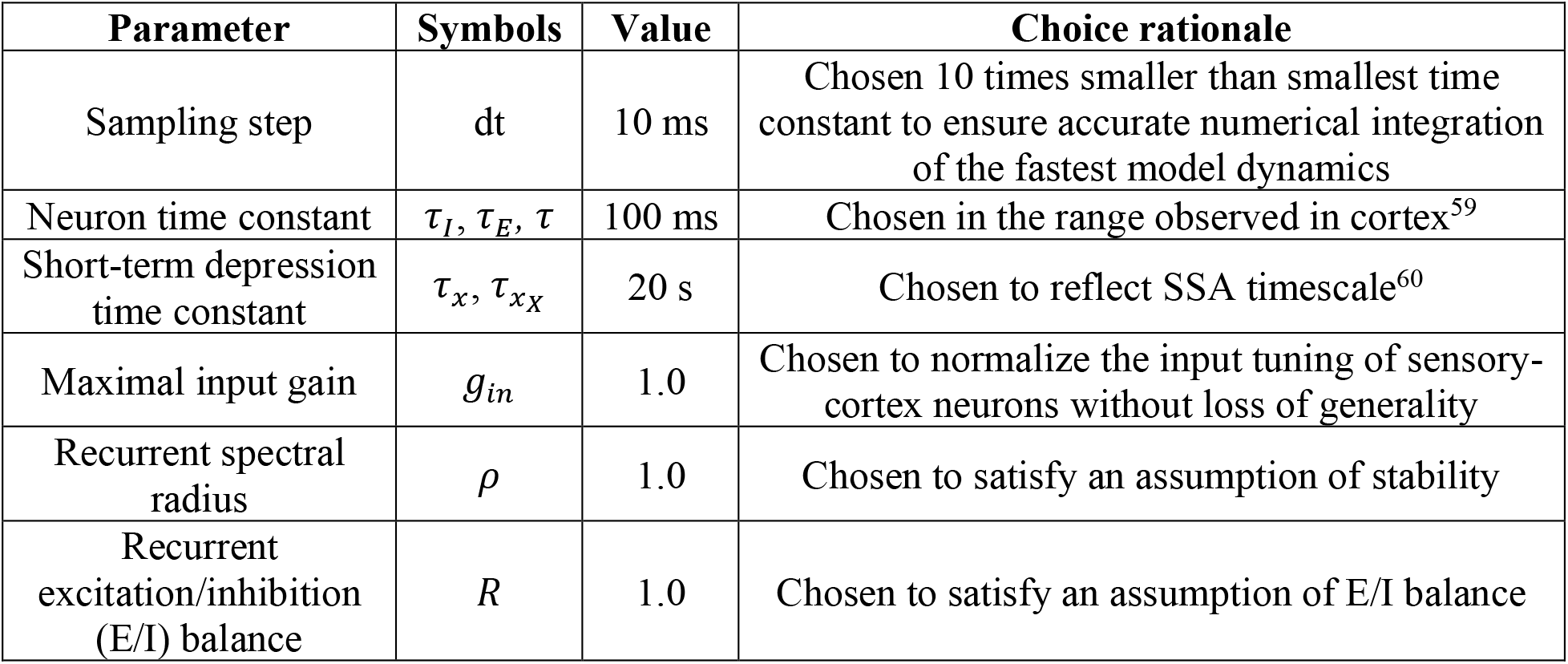
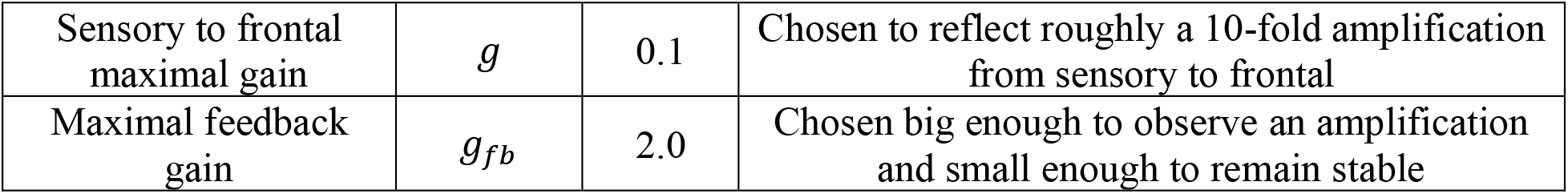
Parameters of models and task. Analysis.

| Parameter | Symbols | Value | Choice rationale |
| --- | --- | --- | --- |
| Sampling step | dt | 10 ms | Chosen 10 times smaller than smallest time constant to ensure accurate numerical integration of the fastest model dynamics |
| Neuron time constant | $\tau_I, \tau_E, \tau$ | 100 ms | Chosen in the range observed in cortex<sup>59</sup> |
| Short-term depression time constant | $\tau_x, \tau_{xX}$ | 20 s | Chosen to reflect SSA timescale<sup>60</sup> |
| Maximal input gain | $g_{in}$ | 1.0 | Chosen to normalize the input tuning of sensory-cortex neurons without loss of generality |
| Recurrent spectral radius | $\rho$ | 1.0 | Chosen to satisfy an assumption of stability |
| Recurrent excitation/inhibition (E/I) balance | $R$ | 1.0 | Chosen to satisfy an assumption of E/I balance |
| Sensory to frontal maximal gain | $g$ | 0.1 | Chosen to reflect roughly a 10-fold amplification from sensory to frontal |
| Maximal feedback gain | $g_{fb}$ | 2.0 | Chosen big enough to observe an amplification and small enough to remain stable |

#### Analysis

##### Dimensionality reduction

To study the underlying low-dimensional representation we used principal component analyses (PCA)^61^. Specifically, we applied PCA separately to the sensory and frontal networks, while either varying only the probability of occurrence of stimuli, or the interstimulus interval.

##### Convex hull ellipse fitting

As neural representations of simulated neural responses in principal components resemble ellipses, we quantitatively measure the distortion caused by frequency of stimuli (probability or interstimulus interval) by fitting ellipses to the convex hull of two principal components of the firing rate response of excitatory neurons. We use only the last 50 trials to allow for time for the short-term depression to produce stable representations. To fit the ellipses we first perform a linear fit:

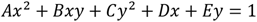

Where *x* and *y* are the coordinates of the convex hull of two principal components, *A*, *B*, *C*, *D* and *E* are the linear weights obtained through least square linear regression.

The we compute the parameters of the ellipses as follow:

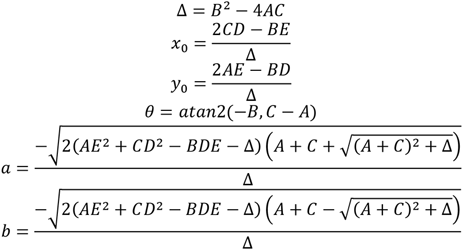

Where *x*_0_ and *y*_0_ are the coordinates of the center of the ellipse, *θ* is the rotation of the ellipse, *a* and *b* are the semi major and minor axes of the ellipse.

When the convex hull is a triangle, the initial ellipse fit occasionally fails — a rare occurrence observed only in the active condition within the frontal region. To address this issue, we introduce isotropic Gaussian noise to the convex hull coordinates only when the initial fit fails. The noise has zero mean and a standard deviation set to 0.01 times the average range of the first and second coordinates.

## Acknowledgement

This work was supported by the National Institutes of Health (NS086085, MH137325), and by the France-Stanford Center for Interdisciplinary Studies.

